# Ultrafast and accurate sequence alignment and clustering of viral genomes

**DOI:** 10.1101/2024.06.27.601020

**Authors:** Andrzej Zielezinski, Adam Gudyś, Jakub Barylski, Krzysztof Siminski, Piotr Rozwalak, Bas E. Dutilh, Sebastian Deorowicz

## Abstract

Viromics produces millions of viral genomes and fragments annually, overwhelming traditional sequence comparison methods. We introduce Vclust, a novel approach that determines average nucleotide identity by Lempel-Ziv parsing and clusters viral genomes with thresholds endorsed by authoritative viral genomics and taxonomy consortia. Vclust demonstrates superior accuracy and efficiency compared to existing tools, clustering millions of virus genomes in a few hours on a mid-range workstation.

## Main text

Metagenomics and viromics are uncovering new viruses at an unprecedented rate, but recognizing which sequences were seen before remains challenging^12^. Calculating average nucleotide identity (ANI), essential for classification, is limited by the scalability of alignment tools like VIRIDIC^3^, which is recommended by the International Committee on Taxonomy of Viruses (ICTV) to calculate intergenomic similarities and delineate bacteriophage species and genera. Large-scale sequence comparisons thus primarily rely on efficient, albeit less accurate *k*-mer approaches such as sketching (FastANI^4^) or sparse approximate alignments (skani^5^). Moreover, most tools lack clustering functionality or do not scale to large metagenomic datasets (Table S1).

Vclust is a fast alignment-based method that calculates ANI variants for complete and fragmented viral genomes and supports clustering cutoffs according to ICTV and Minimum Information about an Uncultivated Virus Genome (MIUViG) standards^23^. It integrates three new components (Figure 1a). First, a new Kmer-db version analyzes all genomic *k*-mers, overcoming sampling limitations inherent in sketching. Second, LZ-ANI efficiently estimates ANI using a novel Lempel-Ziv parsing algorithm (Figure 1b, Methods) with high sensitivity and accuracy. Third, Clusty offers efficient implementation of six clustering algorithms suited for sparse distance matrices with millions of genomes (Figure 1c).

**Figure 1.**
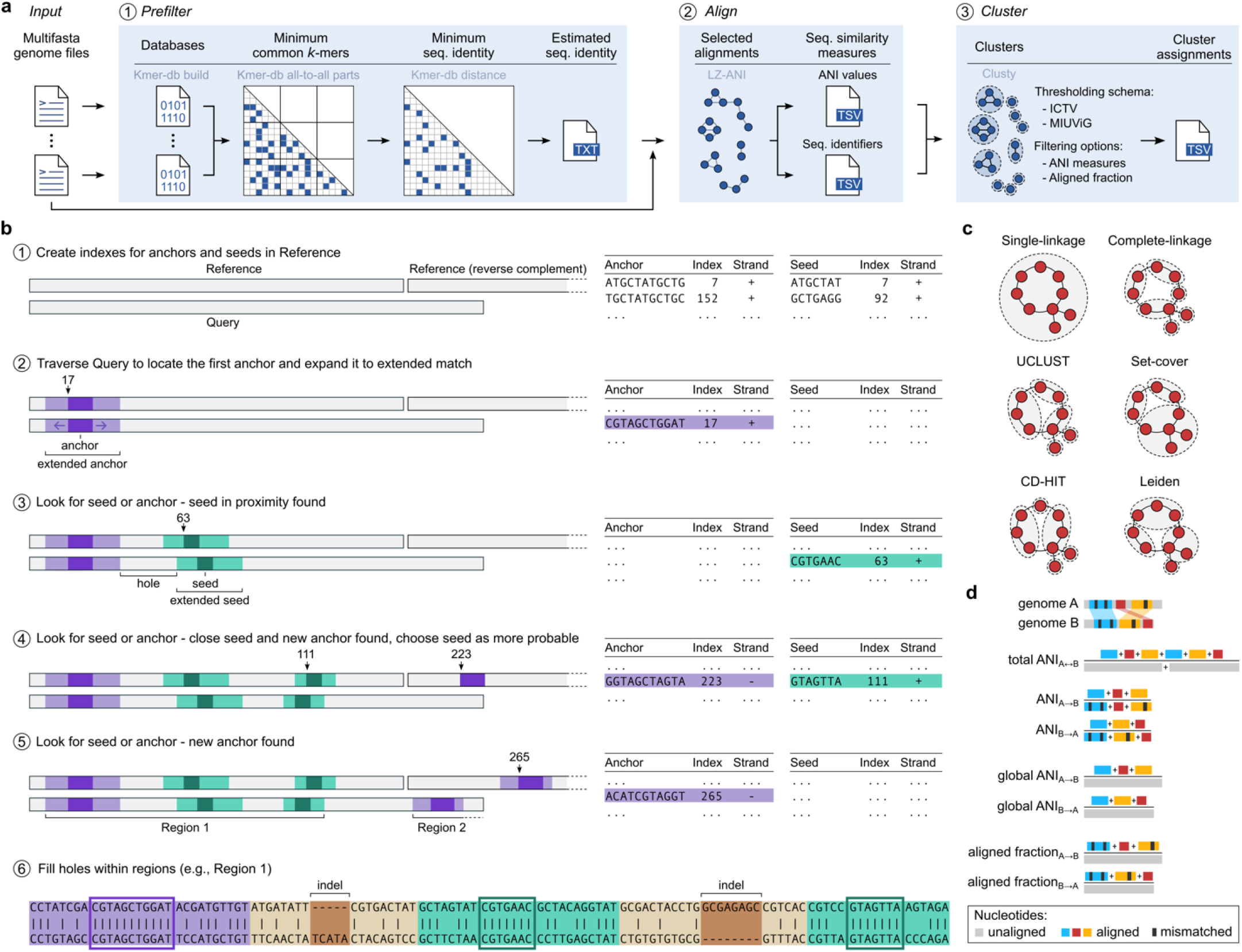
Vclust algorithm and features. **a**, Vclust’s workflow: (1) prefilter similar genome sequence pairs with sufficient *k*-mer-based identity estimated using Kmer-db 2, (2) align similar genome pairs and calculate ANI using LZ-ANI, and (3) cluster genomes based on defined cutoffs using Clusty. **b**, Sequence alignment using Lempel-Ziv parsing (see Methods). **c, d**, Vclust’s clustering algorithms and ANI options.

We first tested Vclust’s accuracy of total ANI (tANI) estimation (Figure 1d) among 10,000 pairs of phage genomes containing simulated mutations, including substitutions, deletions, insertions, inversions, duplications, and translocations (Methods, Table S2). Alignment-based tools, Vclust and VIRIDIC, provided tANI values close to the expected ones, with mean absolute error (*MAE*) of 0.3% and 0.7%, respectively, outperforming FastANI (*MAE* = 6.8%) and skani (21.2%) (Figure 2a). Vclust predictions consistently approached expected values as tANI increased, while VIRIDIC underestimated tANI (Figure 2a). Among genome pairs above the ICTV’s species threshold (tANI ≥95%^3^, *n* = 1,188), Vclust reported only 22 pairs below the threshold, whereas VIRIDIC underestimated nearly ten times more (*n* = 210, Table S3).

**Figure 2.**
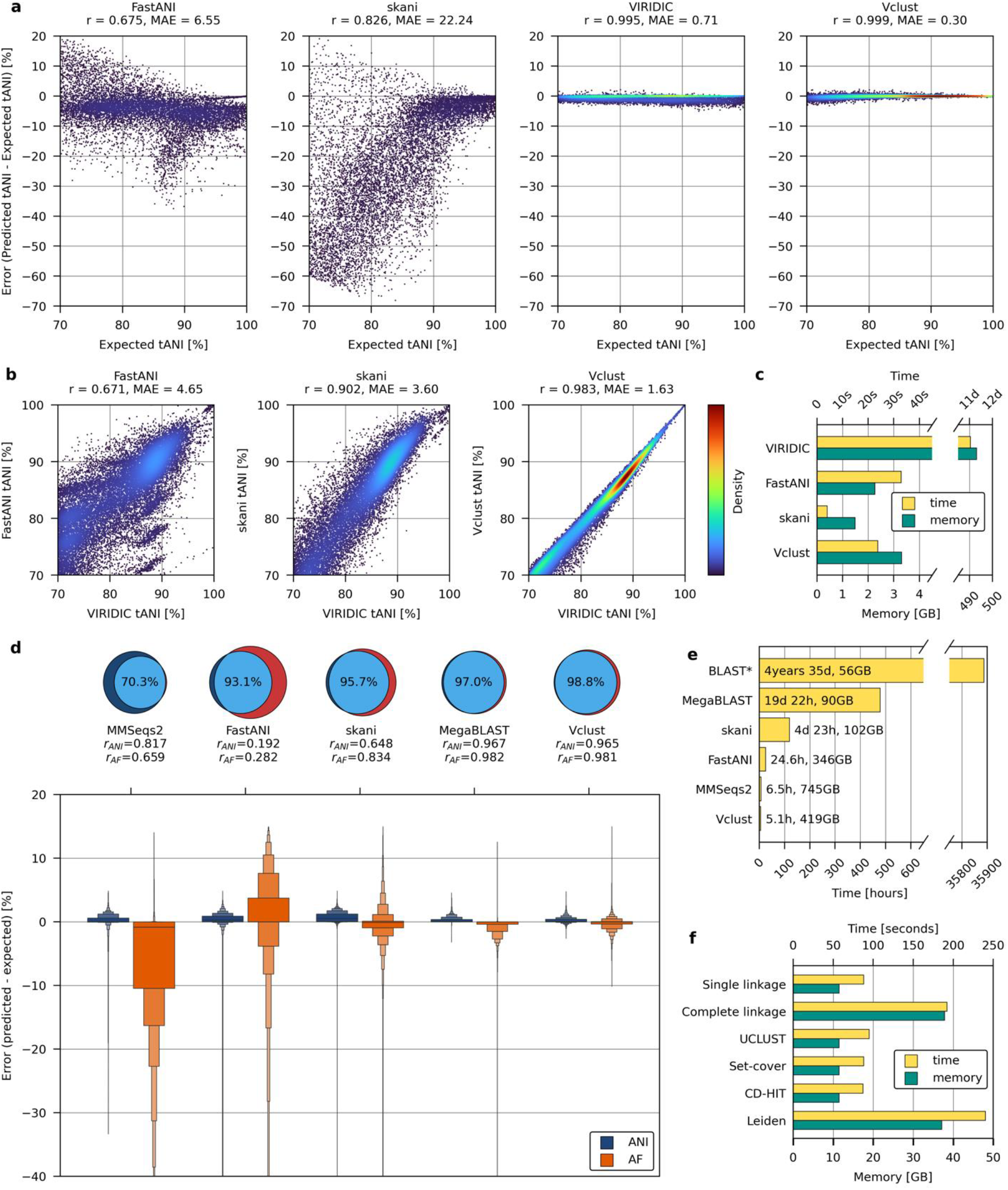
Comparison of Vclust with other tools on various datasets. **a**, Difference between predicted and expected total ANI (tANI) values for 10,000 bacteriophage genome pairs with simulated mutation events. **b**, Correlations with VIRIDIC tANI values for 22,606 complete bacteriophage genome pairs. **c**, Wall time and peak memory usage for processing 4,244 bacteriophage genomes (32 threads). Vclust and VIRIDIC include clustering, while FastANI and skani only calculate ANI. **d**, Venn diagrams comparing numbers of contig pairs meeting MIUViG thresholds (ANI ≥95% and AF ≥85%) predicted by blastn (purple) and other tools (red). The boxen plot shows the error distribution of predicted ANI and AF values relative to corresponding blastn-based reference values for 4,361,743 contig pairs meeting MiUVUG thresholds. The center line denotes the median, while each box level from the median contains half of the remaining observations. **e**, Wall time and peak memory usage for calculating ANI and AF among 15,677,623 IMG/VR contigs (64 threads). blastn values were estimated from a random sample of 1,000 query contigs. **f**, Wall time and peak memory usage of Vclust’s clustering algorithms for grouping IMG/VR contigs into vOTUs.

Next, we determined tANI using VIRIDIC in an all-to-all comparison of 4,244 bacteriophage genomes. Vclust had a higher correlation with VIRIDIC tANI (Pearson’s *r* = 0.983) than skani (*r* = 0.902) and FastANI (*r* = 0.671) across the entire tANI range ≥70% (22,606 genome pairs, Figure 2b) and outperformed both tools within their optimal range ≥80%^4,5^.

We then compared the consistency of the bacteriophage species-level groupings (tANI ≥95%) with the official ICTV taxonomy (Methods). Surprisingly, all tools showed low agreement with ICTV, Vclust showing the highest congruence (73%), followed by VIRIDIC (69%), FastANI (40%), and skani (27%). Upon examining genome pairs where both Vclust and VIRIDIC diverged from the ICTV’s classification, we found inconsistencies in 50 ICTV taxonomic proposals (Tables S4-S5). Excluding these cases improved the agreement of both tools with ICTV taxonomy, with Vclust (95%) retaining superiority over VIRIDIC (88%) and the other tools. Given Vclust’s high agreement with ICTV taxonomy, accurate tANI determination, and processing speed >39,000 times faster than VIRIDIC (Figure 2c, Table S6), it emerges as the prime tool for bacteriophage classification.

We then evaluated Vclust’s performance in clustering virus contigs into operational taxonomic units (vOTUs) based on MIUViG’s thresholds (ANI ≥95% and alignment fraction (AF) ≥85%, Figure 1d). As a reference, we used ANI and AF values determined by blastn^6^ between a sample of over 4 million contig pairs from IMG/VR, which met the MIUViG thresholds (Methods). Vclust recovered the highest number of contig pairs (99%), followed by MegaBLAST (97%), skani (96%), FastANI (96%), and MMseqs2^7^ (70%, Figure 2d). Both Vclust and MegaBLAST produced ANI and AF estimates consistently with the blastn values (Pearson *r* >0.96) outperforming the other tools (*r* = 0.2-0.8). On average, ANI and AF values obtained by Vclust and MegaBLAST showed minimal deviation from the expected values (*MAE* <0.5%, Table S7), with Vclust having the narrowest error range among all the tools (Figure 2d).

We tested the scalability of the tools using the entire IMG/VR database of 15,677,623 virus contigs. Vclust performed sequence identity estimations for ∼123 trillion contig pairs and alignments for ∼1.5 billion pairs, resulting in 5-8 million vOTUs depending on the clustering algorithm (Table S8). Vclust was over 75 times faster than MegaBLAST, >3-fold faster than skani or FastANI, and slightly faster than the fastest mode of MMseqs2 (Figure 2e,f, Table S9). The Vclust vOTUs are generally consistent with those identified by MegaBLAST, with Vclust clustering approximately 75,000 more contigs on average, indicating higher sensitivity (Table S8).

In conclusion, Vclust surpasses the current state-of-the-art methods in viral genome comparison in both accuracy and speed, remaining effective in datasets of millions of sequences. It provides a complete solution for calculating intergenomic similarities and clustering both complete and partial virus genomes using various ANI measures and clustering algorithms. Given the astonishing diversity of viruses in metagenomic data, we believe that Vclust will be essential for large-scale dereplication and taxonomic classification of viral sequences. Finally, for small-sized projects, Vclust is accessible as a web service (http://www.vclust.org/), and the core components Kmer-db, LZ-ANI, and Clusty are available as versatile software tools for independent use in other fields involving large-scale sequence comparisons or clustering.

## Online Methods

### Methods overview

Vclust is a workflow integrating three new tools:

1. Kmer-db 2 that performs initial *k*-mer-based estimation of sequence identity of all genome pairs (section “Sequence identity estimation: Kmer-db 2”).
2. LZ-ANI that aligns sequence pairs with nucleotide identity exceeding a specified threshold and calculates average nucleotide identity (ANI) and aligned fraction (AF) measures (sections “Sequence alignment algorithm: LZ-ANI” and “Calculating ANI and aligned fraction”).
3. Clusty clustering sequences based on ANI and/or AF criteria (section “Clustering sequences: Clusty”).

We implemented Kmer-db 2, LZ-ANI, and Clusty in C++20. Designed as stand-alone tools, they can be easily adapted to various sequence comparison and clustering tasks (see Code availability). Vclust itself is a Python script that integrates these tools to calculate and cluster viral genomic sequences (section “Vclust implementation”).

### Sequence identity estimation: Kmer-db 2

Kmer-db 2 is an updated tool for *k*-mer-based estimation of pairwise similarities among nucleotide sequences that introduces several improvements enabling the processing tens of millions of sequences.

First, unlike its predecessor, which stored similarity values in RAM as a dense matrix^8^, Kmer-db 2 uses sparse matrices that retain only non-zero elements in all-to-all pairwise genome comparison mode (‘all2all-sp’), allowing the algorithm to handle large and diverse genome sets. Second, Kmer-db 2 supports genome datasets partitioned into multiple input files, each generating a separate Kmer-db database. A new mode, ‘all2all-parts’, calculates shared *k*-mers within and across databases, optimizing memory by sequentially loading one or two analyzed databases into RAM. This mitigates memory limitations at the cost of computational overhead due to the repeated database loading. Third, Kmer-db 2 can further minimize RAM usage by storing only genome pairs that meet a minimum threshold of shared *k*-mers and sequence identity. Finally, all modes in Kmer-db 2 support multithreading, except for the distance calculation step, which is fast enough to not require parallelization.

### Sequence alignment algorithm: LZ-ANI

The LZ-ANI algorithm uses Lempel-Ziv (LZ) parsing^9^ to align two sequences (the query and the reference).

First, the algorithm constructs two indices (dictionaries): for anchors and seeds. The anchor index maps all *a*-mers (substrings of length *a*) from both strands of the reference sequence to their positions, while the seed index performs the same mapping for shorter *s*-mers (Figure 1b step 1).

Next, the query is read from left to right using a sliding window of *a* nucleotides, moving one nucleotide at a time. The parsed *a*-mers are used to search the anchor index for matches in the reference. Upon finding an exact match, the algorithm extends it in both directions (Figure 1b step 2). In each direction, a window of size *aw* slides until it encounters more than a certain number of mismatches (*am)* at a time. Then, the extensions of terminal windows are trimmed to remove poorly aligned ends until they have at least *ar* exactly matched nucleotides. This extended anchor initiates the first *region*, which corresponds to a local alignment, and is constructed as described below.

The algorithm then moves to the next nucleotide after the extended anchor and looks for *a*-mers (anywhere in the reference) and *s*-mers (within *r* nucleotides from the end of the extended match in the reference) in the dictionaries. Four scenarios may arise:

1. No anchor or seed is found: shift by one position in the query and repeat the process of finding a new anchor or a seed match. However, if the distance in the query between the current position and the end of the previous match exceeds *q* nucleotides, the seed search is discontinued.
2. Only a seed match is found: extend the seed similarly to the initial anchor match, append it to the region, and continue the search for a new anchor or seed match (Figure 1b step 3).
3. Only an anchor match is found: close the current region and extend the anchor match to initiate a new region (Figure 1b step 5).
4. Both anchor and seed matches are found: select the match less likely to occur by chance, based on their lengths, seed proximity (*r* nucleotides), and the reference sequence length, leading to either scenario 2 or 3 (Figure 1b step 4).

Upon closing a *region*, the algorithm realigns the nucleotide stretches between all the extended matches within the region (Figure 1b step 6). This realignment aims to maximize the number of matching nucleotides between neighboring extended matches by allowing for a single multi-symbol insertion in the reference or query sequence. As a result, the region represents a local alignment containing both matched and mismatched nucleotides, along with approximated indel fragments. To remove spurious alignments, regions shorter than *g* nucleotides are excluded from further analysis.

All LZ-ANI parameters are adjustable and were optimized for virus genome sequences. Optimization was conducted using a dataset comprising 10,000 pairs of complete genomes with simulated mutations, encompassing various levels of substitution, insertion, deletion, duplication, inversion, and translocation events. Detailed parameter settings are provided in Table 1.

**Table1.** Parameters of LZ-ANI sequence aligner. All length values are given in nucleotides.

| Parameter | Role | Default value |
| --- | --- | --- |
| $a$ | anchor length | 11 |
| $s$ | seed length | 7 |
| $aw$ | window length for match (anchor or seed) extension | 15 |
| $am$ | maximum number of mismatches in the extension window | 7 |
| *ar* | minimum number of consecutive matches at the end of the extension | 3 |
| *r* | maximum distance between approx. matches in reference | 35 |
| *q* | maximum dist. between approx. matches in reference | 35 |
| *g* | minimum region length to consider | 40 |

The LZ-ANI tool reads input sequences and stores them in RAM using a compact format, with two nucleotides represented per byte. The tool processes sequences in parallel, with each thread comparing a reference sequence to all other sequences. By default, the tool performs all-versus-all pairwise alignments but it can accept a filter containing a list of sequence pairs to align, e.g., a file obtained by Kmer-db (used by Vclust by default).

### Calculating ANI and aligned fraction

LZ-ANI alignment between query (*A*) and reference (*B*) allows direct calculation of:

- *L(A, B)* - the total length of all regions when aligning query *A* to reference B,
- *M(A, B) -* the total number of matching nucleotides in all regions,
- *N(A, B)* - the number of regions.

These values are used to compute seven sequence similarity measures, as follows:

- ANI for *A* and *B*: 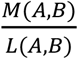
- ANI for *B* and *A*: 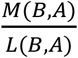
- Aligned fraction (AF) of query *A* to reference *B*: 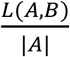
- Aligned fraction (AF) of query *B* to reference A: 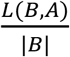
- Global ANI for *A* and *B*: 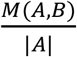
- Global ANI for *B* and *A*: 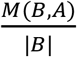
- Total ANI: 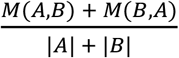.

### Clustering sequences: Clusty

Clusty is a versatile package facilitating rapid clustering across diverse data types, using six algorithms: single linkage, complete linkage, UCLUST^10^, greedy set cover^7^, CD-HIT^11^, and Leiden^12^. Our implementations of these algorithms have been optimized for sparse distance matrices. A linear memory complexity with the number of distances allows clustering tens of millions of objects, provided the matrix remains sufficiently sparse.

Clusty uses threshold-based clustering, assigning an object to a cluster if its distance from the cluster does not exceed a user-defined threshold. This distance criterion varies based on the algorithm and can refer to the closest cluster member, the furthest member, or the centroid. While UCLUST, greedy set cover, and CD-HIT are inherently threshold-based algorithms, single and complete linkage algorithms construct dendrograms that can be pruned at customizable distance thresholds. As Clusty uses sparse data representation, which contains only non-zero values, it assumes that all the input values meet the distance or similarity threshold. However, our package allows for clustering data at more stringent thresholds through additional filtering of any combinations of distance/similarity values (e.g., tANI, ANI, AF) and/or other measure values (e.g., minimum/maximum number of alignments, minimum/maximum number of matched nucleotides). Consequently, the matrix provided to Clusty does not need to be sparse; the tool can handle dense matrices and apply filtering at the loading stage.

Clusty interprets input data as a graph, with vertices representing objects and edges representing connections between them. Details on the clustering algorithms provided by Clusty, including their time complexities, are given in Table 2.

**Table 2.** Clustering algorithms available in Clusty, with *n* representing the number of objects (vertices) and *e* the number of distances (edges) in the data set (graph).

| Algorithm | Details | Time complexity |
| --- | --- | --- |
| Single linkage | Hierarchical agglomerative clustering with a distance between groups defined as a distance between their closest members. Equivalent to finding all consistent subgraphs in a graph. Performed using breadth-first search. | $O(e)$ |
| Complete linkage | Hierarchical agglomerative clustering with a distance between groups defined as a distance between their furthest members. Equivalent to finding a disjoint set of complete subgraphs covering the entire graph. Fast identification of clusters to merge is performed by storing distances in a heap. | $O(e \log e)$ |
| UCLUST | Greedy clustering with objects investigated from the most to the least representative (in a sequence domain representativeness usually translates to length). The first object (e.g. longest sequence) becomes a centroid; the following objects are either (a) assigned to the closest of centroids they are connected with or (b) become new centroids if they are not connected to any of the existing ones. | $O(e)$ |
| Greedy set cover | Greedy clustering with objects investigated in descending order with respect to the number of neighbors. Every unassigned object becomes a new centroid with all connected objects being | $O(n \log n + e)$ |
|  | assigned to it. Equivalent to MMseqs2 mode 0 clustering. |  |
| CD-HIT | Greedy clustering with objects investigated from the most to the least representative (in a sequence domain representativeness usually translates to length). Every unassigned object becomes a new centroid with all connected objects being assigned to it. Equivalent to MMseqs mode 2 clustering. | $O(e)$ |
| Leiden | Iterative heuristic for finding communities in networks. It consists of three phases: (1) local moving of nodes, (2) refinement of the partition, and (3) aggregation of the network using the refined and the non-refined partitions. | unknown<br>(provided by *igraph* library<sup>13</sup>) |

### Vclust implementation

Vclust is a Python tool integrating Kmer-db 2, LZ-ANI, and Clusty for streamlined computation of intergenomic sequence similarities and clustering of viral genomes. Vclust implements three commands: prefilter, align, and cluster (Figure 1a). Prefilter and align accept a single FASTA file containing viral genomic sequences or a directory of FASTA files (one genome per file), with support for gzipped inputs and outputs.

The prefilter command uses Kmer-db 2 to establish a pre-alignment filter, which screens out dissimilar genome pairs before conducting pairwise alignments. This process reduces the number of potential genome pairs to only those with sufficient *k*-mer-based sequence similarity (i.e., minimum number of common *k*-mers and/or the minimum sequence identity of the shorter sequence). To reduce memory usage, large sets of genome sequences can be processed in smaller, equally-sized batches using the built-in multi-fasta-split C++ tool.

The align command uses LZ-ANI to perform pairwise sequence alignments and compute ANI and AF measures between genome pairs identified by the pre-alignment filter. If the filter is not provided, Vclust aligns all possible genome pairs. Output consists of two TSV files: one containing ANI measures for genome pairs and the other listing genome identifiers sorted by decreasing sequence length. Both TSV files serve as inputs for Vclust’s clustering.

The cluster command uses Clusty for genome clustering, allowing users to specify a similarity measure (e.g., tANI, ANI) and its threshold for clustering genomes, with optional additional filtering thresholds for other similarity measures, including aligned fraction. Output includes a TSV file listing genome identifiers and numerical cluster identifiers (including identifiers for singleton genomes). Alternatively, Vclust can output representative genomes instead of numerical cluster identifiers, particularly useful for dereplication tasks.

Vclust is primarily built for dereplication and clustering across a range of sequence identity values. Computational performance may decline with very large datasets of highly similar genomes (e.g., tens of thousands of sequences from the same species) as after prefiltering, many sequence pairs remain for alignment and clustering. While users may adjust the cutoff for triggering an alignment to address this issue, future work will focus on developing methods for handling large homogeneous datasets, including bacterial genomes.

### Benchmarking

#### Running time

All runtimes were benchmarked on an AMD Epyc 9554 CPU clocked at 3.1 GHz machine with 64 cores and 1152 GB RAM. Unless otherwise specified, all tools were run using 64 threads. The exact commands are shown in Tables S6 and S9.

#### Evaluating tANI accuracy

The tANI accuracy of Vclust v1.0, FastANI v1.33^4^, skani v0.2.1^5^, and VIRIDIC v1.1^3^ was assessed using two reference sets. The first set comprised 22,606 reference tANI values ranging from 70% to 100%, as determined by VIRIDIC across 4,244 complete genomes of bacteriophages affiliated with the International Committee on Taxonomy of Viruses (ICTV). Since FastANI and skani do not directly report tANI, their values were calculated from ANI, AF, and genome lengths: *tANI* = (*ANI*_1_ × *AF*_1_ × *LEN*_1_ + *ANI*_2_ × *AF*_2_ × *LEN*_2_)/ (*LEN*_1_ + *LEN*_2_). The second reference set contained expected (true) tANI values in the 70-100% range, derived from 10,000 pairs of bacteriophage genomes subjected to simulated mutations, including different levels of substitution, insertion, deletion, duplication, inversion, and translocation events. Specifically, we randomly selected 100 genomes from the bacteriophage dataset and generated 100 copies of each genome. For each genome copy, we introduced mutations using Mutation-Simulator v3.0.2^14^ by randomly selecting a combination of mutation events and their corresponding frequencies (Table S5). The expected (true) tANI between each copy and reference genome was determined based on the Variant Call Format (VCF) file produced by Mutation-Simulator, describing the exact locations of introduced mutations and the number of altered nucleotides.

#### Evaluating ANI and AF accuracy

The ANI and AF values predicted by Vclust, FastANI, skani, MegBLAST v2.13.0+, and MMseqs2 v2fad714b525f1975b62c2d2b5aff28274ad57466^7^ were compared to reference ANI and AF values determined by blastn 2.13.0+^6^. Since running blastn on the entire IMG/VR v4.1 database was not feasible, we subsampled 94,225 viral contigs and performed an all-to-all blastn search to identify 4,361,743 contig pairs meeting the MIUViG thresholds (ANI ≥ 95% and AF ≥ 85%). MegaBLAST, MMseqs, and blastn outputs were used by anicalc script from CheckV v1.0.3^15^ to compute ANI and AF values. Pearson correlation and mean absolute error (MAE) between the predicted and expected ANI and AF values were calculated based on the 4,361,743 contig pairs meeting MIUViG thresholds (ANI ≥ 95% and AF ≥ 85%) determined by blastn. Given the high level of sequence identity of the reference contig pairs, if a tool did not return a result for a given contig pair, the ANI and AF values were set to zero for that pair.

#### Evaluating clusterings

The agreement between clustering results from different tools and the reference clustering was assessed using the Adjusted Rand Index (ARI). ARI assesses clustering similarity by comparing the number of correct clustering overlaps and disagreements^16^ against those expected by chance. An ARI of 0 indicates random assignment, while a score of 1 indicates a perfect match. We used the scikit-learn v1.3.2^17^ implementation of the ARI.

## Data Availability

The dataset of complete genomes of 4,244 bacteriophages is available at https://doi.org/10.6084/m9.figshare.25907833.v1. The dataset including 10,000 bacteriophage genome sequences containing simulated mutations and corresponding expected total ANI values is available at https://doi.org/10.6084/m9.figshare.25907815.v1. The dataset of 94,225 metagenomic viral contigs sampled from IMG/VR v4.1 with the expected blastn-based ANI and AF values is available at https://doi.org/10.6084/m9.figshare.25907803.v1. The full dataset of 15,677,623 contigs from IMG/VR v4.1 is available at https://genome.jgi.doe.gov/portal/IMG_VR/. Source data are provided with this paper.

## Code Availability

Vclust is available as a standalone tool at https://github.com/refresh-bio/vclust and as a web service at http://www.vclust.org. Kmer-db 2, LZ-ANI, and Clusty are available respectively at https://github.com/refresh-bio/kmer-db, https://github.com/refresh-bio/LZ-ANI, and https://github.com/refresh-bio/clusty.

## Acknowledgments

This work is supported by the National Science Centre, Poland, project DEC-2022/45/B/ST6/03032 (to AG and SD), the European Research Council (ERC) Consolidator grant 865694: DiversiPHI, the Deutsche Forschungsgemeinschaft (DFG, German Research Foundation) under Germany’s Excellence Strategy – EXC 2051 – Project-ID 39071386 (to BED), the European Union’s Horizon 2020 research and innovation program, under the Marie Skłodowska-Curie Actions Innovative Training Networks grant agreement no. 955974 (VIROINF) (to BED), the Alexander von Humboldt Foundation in the context of an Alexander von Humboldt-Professorship founded by German Federal Ministry of Education and Research (to BED and PR), and the Polish Ministry of Science and Higher Education under the programme “Perły Nauki”, project number PN/01/0063/2022 (to PR). The computations were partially performed at the Poznan Supercomputing and Networking Center (grant numbers pl0243-01 and pl0074-02).

## Author Contributions Statement

AZ, AG, JB, and SD designed the study. AZ conducted the comparative analyses and developed the web service. AG designed and developed Clusty with input from KS and SD. AG and SD designed and developed Kmer-db. SD designed and developed LZ-ANI. AZ, AG, and SD developed the Vclust tool. AZ and JB compared predictions to the ICTV taxonomy and reviewed ICTV proposals. AZ, JB, AG, PR, BED, and SD analyzed the results. AZ and AG designed figures, with inputs from SD, JB, KS, and BED. AZ, AG, and SD wrote the manuscript with substantial contributions from JB, PR, and BED. All authors reviewed and approved the manuscript.

